# Live-cell STED microscopy of mitochondrial cristae

**DOI:** 10.1101/640920

**Authors:** Till Stephan, Axel Roesch, Dietmar Riedel, Stefan Jakobs

## Abstract

Mitochondria are highly dynamic organelles that exhibit a complex inner architecture. They exhibit a smooth outer membrane and a highly convoluted inner membrane that forms invaginations called cristae. Imaging cristae in living cells poses a formidable challenge for light microscopy. Relying on a cell line stably expressing the mitochondrial protein Cox8A fused to the SNAP-tag and using STED (*st*imulated *e*mission *d*epletion) super-resolution microscopy, we demonstrate the visualization of cristae dynamics in cultivated human cells.

## Introduction

Mitochondria form tubular and highly dynamic networks in mammalian cells that constantly undergo fusion and fission events^1,2^. They are double-membrane organelles that exhibit a smooth outer membrane and a highly convoluted inner membrane. Cristae are invaginations of the inner membrane that generally adopt tubular or lamellar shapes and project into the matrix space. The cristae architecture adapts to different cellular conditions and is changed upon various processes^3,4^. However, the actual cristae dynamics in these processes are poorly understood.

A challenge for studying cristae dynamics is the small size of mitochondria. The diameter of mitochondrial tubules is generally between 200 and 700 nm, and in many mammalian cell types the crista-to-crista distance is below 100 nm. Hence, because of the diffraction limit in optical microscopy (∼250 nm, depending on the wavelength), visualization of cristae dynamics has been a notorious challenge^5^.

Electron microscopy provides the required resolution, but it is restricted to fixed samples. So far, the overall mitochondrial dynamics and the cristae movements, difficulties in labelling and concerns on light induced photo-toxicity have hampered the visualization of single cristae dynamics using live-cell nanoscopy^6,7^. In fixed cells, cristae were first visualized using isoSTED nanoscopy ^8^. Using structured illumination microscopy providing a resolution of around 100 nm, mitochondrial substructures were resolved in living cells^9-11^. However, the discrimination between individual cristae and groups of cristae remained difficult because of the limited resolution.

We overcome this problem by using diffraction-unlimited STED super-resolution microscopy (STED nanoscopy) in combination with a genome edited human cell line that enables labeling of the cristae with a silicon rhodamine dye. Thereby, we unequivocally visualize the dynamics of individual cristae using nanoscopy.

## Results

### Visualization of the cristae architecture in living cells

We have generated a human HeLa cell line that stably expresses the full length Cytochrome *C* Oxidase Subunit 8A (Cox8A), an integral protein of the mitochondrial inner membrane, fused to the SNAP-tag^12^. As Cox8A is a subunit of complex IV of the respiratory chain, it is expected to be preferentially localized in the crista membrane^13,14^. The fusion construct was targeted to the chromosomal AAVS1 (*a*deno-*a*ssociated *v*irus integration *s*ite 1) Safe Harbor Locus using a CRISPR/Cas9 based genome editing strategy, ensuring largely constant expression levels^15,16^.

Labeling of living cells by adding the cell permeant dye SNAP-Cell SiR to the medium^17^ resulted in brightly fluorescent mitochondria. As expected, diffraction-limited confocal recordings did not reveal any sub-mitochondrial structures, whereas with STED microscopy (excitation: 640 nm, STED: 775 nm), we were able to record cristae in mitochondria of living HeLa cells with ∼50 nm resolution (Fig. 1). Individual cristae could be resolved, typically exhibiting a crista-to-crista distance between 70 and 90 nm, which is fully in line with electron micrographs recorded from the same cell line (Fig.1, Fig. 2A). We were able to record entire cells, thereby getting an overview on the cristae architecture of mitochondria with different shapes or with different subcellular positions (Fig. 1).

**Fig. 1.**
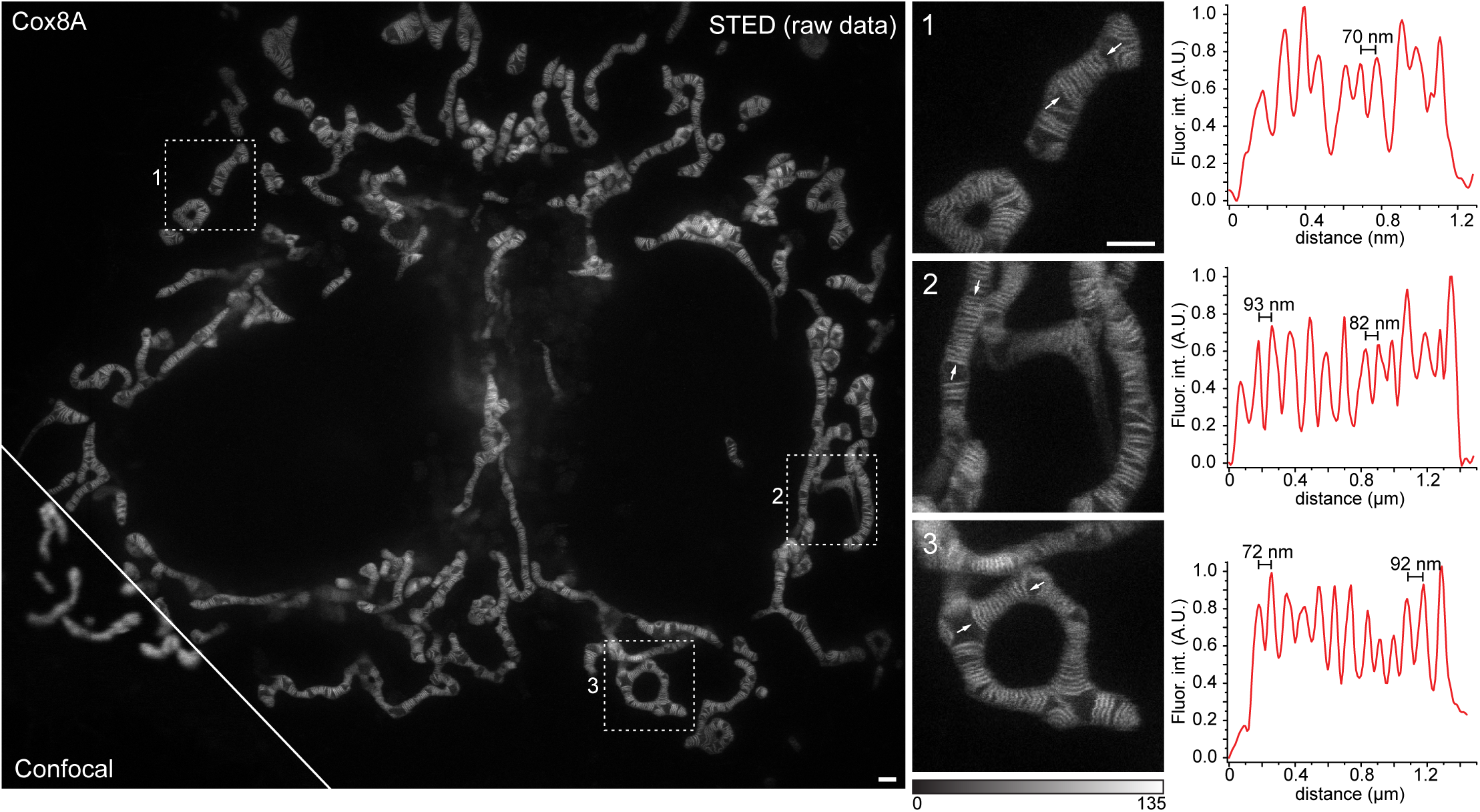
Live-cell Stimulated Emission Depletion (STED) nanoscopy of mitochondrial cristae in HeLa cells. HeLa cells stably expressing Cox8A-SNAP fusion proteins were labelled using SNAP-Cell SiR and visualized with STED nanoscopy. Left: Overview of HeLa cells. Shown is a comparison between confocal and STED resolution. Middle: Magnifications of the areas indicated in the overview image. Right: Fluorescence intensity line profiles measured as indicated in the magnifications. Images show unprocessed raw data without background subtraction. Scale bars: 1 µm.

**Fig. 2.**
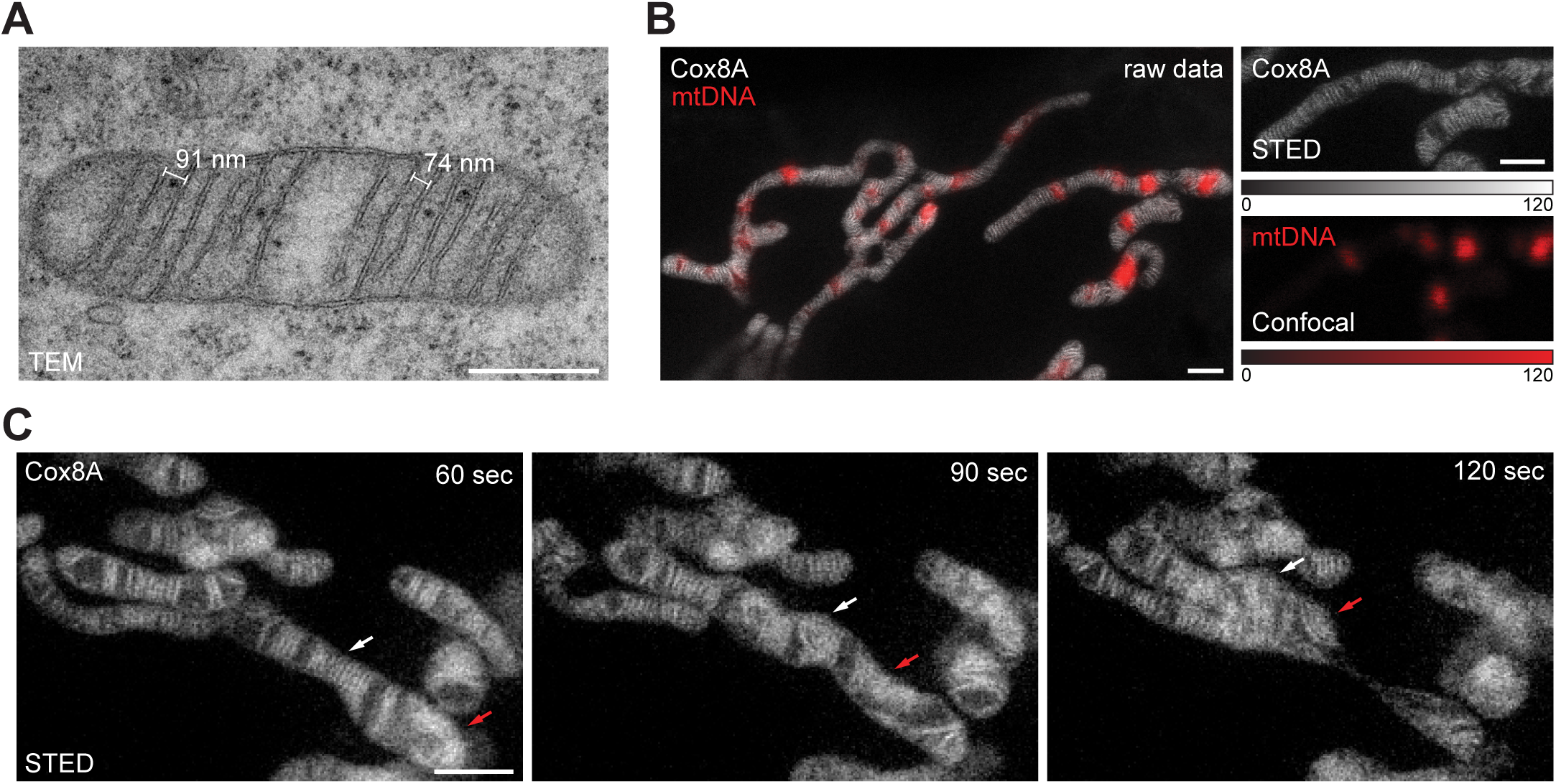
Dynamics of mitochondrial cristae. **A.** Transmission electron microscopy (TEM) of mitochondria from HeLa cells. **B.** Dual-color live-cell imaging of mitochondria. HeLa cells stably expressing Cox8A-SNAP fusion proteins were labelled using SNAP-Cell SiR. In addition to the cristae, mitochondria were labeled for mitochondrial DNA with PicoGreen. The cristae were visualized by STED nanoscopy, the mitochondrial DNA by confocal scanning light microscopy. **C.** Live-cell time-lapse STED nanoscopy of mitochondria. Mitochondria were recorded every 15 seconds. Shown are three frames out of ten. Arrows indicate the behavior of two separated groups of cristae during the fission of a mitochondrion. B. shows unprocessed raw data without background subtraction; in C. photobleaching was compensated by adapting the color table. Scale bars: A: 0.5 µm; B,C: 1 µm.

### Mitochondrial nucleoids occupy the voids between groups of cristae

The cristae frequently occurred in groups, separated by voids of several hundred nanometers size. Labeling of the mitochondrial DNA with the cell-permeable dye PicoGreen allowed us to analyze the location of mitochondrial DNA with respect to the cristae in live mitochondria. Unexpectedly, we found that in this cell line most free spaces in the mitochondrial matrix that were devoid of cristae were occupied by mitochondrial nucleoids (Fig. 2B). We note that mitochondrial nucleoids, because of their low contrast, are invisible in conventional EM and hence the detailed localization of nucleoids has not been shown previously.

### Time lapse imaging of cristae dynamics

Using our labeling strategy, image sequences with 10-20 frames could be recorded. This allowed us to capture individual cristae and their dynamics across larger fields of view. By this we could, for example, capture the dynamics of cristae apparently moving in groups during the fission of a mitochondrial tubule (Fig. 2C; Suppl. Movie 1). Due to the flexibility of the beam-scanning STED imaging scheme, the size of the recorded fields of view can be adapted, facilitating video sequences with frame rates in the second range. Even at this temporal resolution we observed an unexpected level of cristae movements (Suppl. Movie 2 and 3).

## Discussion

The intricate mitochondrial membrane architecture is vital for the functioning of mitochondria as cellular powerhouses. It is widely accepted that the mitochondrial cristae are dynamic structures that are remodeled upon various cellular stimuli, but also upon apoptosis and during ageing^4,18-20^. The studies demonstrating cristae adaptations relied on electron microscopy that can visualize the membrane architecture of mitochondria in fixed cells. However, very little is known about the actual dynamics of these processes, as a clear visualization of the cristae structure in living cells has been a notorious challenge in the past.

The mechanisms of mitochondrial cristae biogenesis are debated and different mechanisms for this process have been suggested^21-28^. To better analyze the formation of cristae and their structure and dynamics under different cellular conditions, a new approach for high-resolution live cell imaging of mitochondrial cristae is urgently needed. We suggest that the Cox8A-SNAP cell line in combination with nanoscopy will be a valuable resource to study cristae dynamics and cristae biogenesis. We note that our images have been recorded with a commercial STED microscope which will allow also non-specialized laboratories to record cristae dynamics. The SNAP-tag allows to change the fluorophore according to the requirements of any light microscopy technique. This versatility will facilitate an immediate comparison of different fluorophores, labelling strategies and imaging modalities on a dynamic and challenging live cell sample. Moreover, the chemical fixation of the double membraned mitochondria is often difficult and we expect this cell line will be a valuable resource for evaluating and systematically improving fixation conditions for optical microscopy. The cell line will be freely available to the scientific community after publication of the manuscript in a peer-reviewed journal.

## Acknowledgements

This work was supported by the Deutsche Forschungsgemeinschaft (DFG, German Research Foundation) funded SFB1286 (project A05) and under Germany’s Excellence Strategy - EXC 2067/1- 390729940 (both to SJ).

## Author contributions

TS and SJ conceived the project; TS, AR and DR performed research; TS and AR analyzed data. SJ wrote the text with comments from all authors.

## Material and Methods

### Cloning of plasmids

To generate the donor plasmid AAVS1-Blasticidin-CAG-Cox8A-SNAP, the plasmid AAVS1-Basticidin-CAG-Flpe-ERT2 was linearized by using the restriction endonucleases *Sal*I and *EcoR*V. Cox8A-SNAP was amplified by PCR from pSNAPf-Cox8A (New England Biolabs, Ipswich, MA, USA) and subsequently integrated into the linearized plasmid by Gibson assembly. AAVS1-Blasticidin-CAG-Flpe-ERT2 was a gift from Su-Chun Zhang (Addgene plasmid #68461; http://n2t.net/addgene:68461; RRID:Addgene_68461). Oligonucleotides for donor Plasmid:

Cox8A-fwd: TCTCATCATTTTGGCAAAGAATTCGTCGACGCCGCCACCATGTCCGTCCTGACGCCG

Cox8A-rev: GAGGTTGATTATCGATAAGCTTGATATCTTAATTAACCTCGAGTTTAAACGCGGATC

The gRNA plasmid PX458-AAVS1 was derived from PX458. In brief, oligonucleotides were annealed by primer annealing and integrated into PX458 after linearization with the *Bbs*I restriction endonuclease. pSpCas9(BB)-2A-GFP (PX458) was a gift from Feng Zhang (Addgene plasmid #48138; http://n2t.net/addgene:48138; RRID:Addgene_48138).

Oligonucleotides for gRNA plasmid:

AAVS1-gRNA-fw: CACCGTGTCCCTAGTGGCCCCACTG AAVS1-gRNA-rev: AAACCAGTGGGGCCACTAGGGACAC

### Cell culture

HeLa cells were cultured in DMEM containing 4,5 g/L Glucose and GlutaMAX(tm) additive (Thermo Fisher Scientific) supplemented with 100 U/ml penicillin and 100 µg/mL streptomycin (Merck Millipore, Burlington, MA, USA), 1mM sodium pyruvate (Sigma Aldrich) and 10% (v/v) fetal bovine serum (Merck Millipore).

### Generation of a stable cell line

To generate the stable cell line, HeLa cells were co-transfected with the plasmids AAVS1-Blasticidin-CAG-Cox8A-SNAP and PX458-AAVS1 using the jetPRIME transfection reagent (Polyplus, Illkirch, France). Starting two days after transfection, the cells were selected using DMEM containing 10 µg/ml Blasticidin S (Invivogen, Toulouse, France) for 7 days. Two weeks after transfection, the cells were stained with DMEM containing 1 µM SNAP-Cell SiR (New England Biolabs, Ipswich, MA, USA) for about 10 minutes. After two washing steps with DMEM (15 min), the cells were detached. Using fluorescence-activated cell sorting (FACS), single cells were transferred into 96 well plates. After about 3 weeks, single cell clones were again stained using the SNAP-cell SiR dye and the expression of Cox8A-SNAP was controlled using confocal fluorescence microscopy using a Leica TCS SP8 (Leica Microsystems, Wetzlar, Germany).

### Staining of live cells for STED nanoscopy

Cells were seeded in glass bottom dishes (ibidi GmbH, Martinsried, Germany) one day before the measurements. Cells were stained with DMEM containing 1 µM SNAP-Cell SiR (New England Biolabs) and optionally 0.1 % (v/v) Quant-IT PicoGreen dsDNA reagent (Thermo Fisher Scientific, Waltham, MA, USA) (15 min, 37 °C). After removing the staining solution and two washing steps with DMEM, the cells were left in the incubator in DMEM for 15-30 minutes to remove unbound dye. Prior to imaging, DMEM was replaced with life cell imaging solution (Thermo Fisher Scientific).

### STED nanoscopy of live cells

STED nanoscopy was performed using a quad scanning STED microscope (Abberior Instruments, Göttingen, Germany) equipped with a UPlanSApo 100x/1,40 Oil objective (Olympus, Tokyo, Japan). The pinhole was set to 0.7 – 1.0 Airy units. A pixel size of 20-25 nm was used. SNAP-Cell SiR was excited at 640 nm and STED was performed at 775 nm wavelength. PicoGreen was excited at 485 nm wavelength. The fluorescence signal was detected using avalanche photo diodes with bandpass filters. For STED imaging of SNAP-Cell SiR, a gating of 0.75-8 ns was applied. Generally, dwell times of 7-10 µs were used. For STED images, each line was scanned 4 to 8 times and the signal was accumulated. For the confocal images, each line was scanned once.

### Image processing

No deconvolution was used. For still images (Fig. 1A, B), unprocessed raw data without background subtraction are shown. For time lapse recordings, photobleaching was compensated using the Bleach Correction function (histogram matching) in the Fiji software.

### Transmission electron microscopy of HeLa cells

HeLa cells were grown on aclar discs (Plano, Wetzlar, Germany) to a confluency of about 60 % and fixed with pre-warmed 2.5 % glutaraldehyde in 0.1 M cacodylate buffer (pH 7.4, 1h, RT). Samples were stored in the fixative over night to complete fixation (4 °C). Cells were washed three times with 0.1M cacodylate buffer and were stained in 1 % (w/v) osmium tetroxide in 0.1 M cacodylate buffer (pH 7.4, 1h, RT). The samples were washed with distilled water (three times, each for 5 min) and stained en-bloc for 30 min in aqueous 1 % (w/v) uranyl acetate (RT, in the dark). Dehydration was performed using an ethanol series of 30, 50, 70 and 100 % (three times, each for 5 min) with a final dehydration step in propylene oxide (5 min). Cells were embedded in Agar 100 epoxy resin. Sections of 50 nm thickness were recorded on a Philips CM120 transmission electron microscope with a TVIPS 2k × 2k slow-scan CCD camera.

## Supplementary Information

### Movie 1 to 3

#### Live-cell time-lapse STED nanoscopy of mitochondria

HeLa cells stably expressing Cox8A-SNAP fusion proteins were labelled using SNAP-Cell SiR and visualized with STED nanoscopy. Mitochondria were recorded every 15 (Movie 1), 10 (Movie 2) or 5 (Movie 3) seconds. Scale bar: 1 µm. The movies show raw data; photobleaching was compensated by adapting the color table. Scale bar: 1 µm.

