## Supplementary figures and images for "Live-cell STED microscopy of mitochondrial cristae"

### Supplemental Movie 1

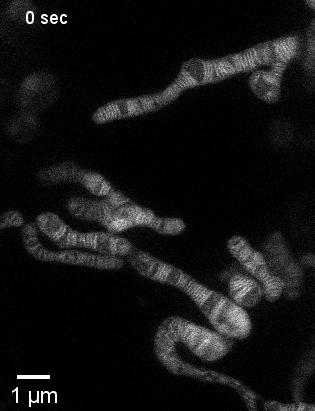

### Supplemental Movie 2

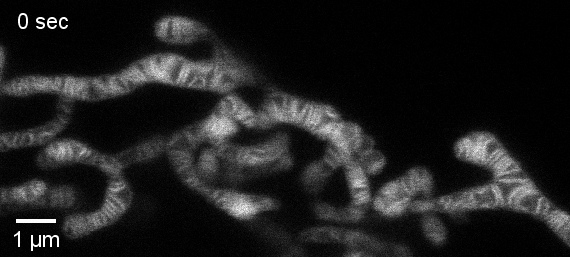

### Supplemental Movie 3

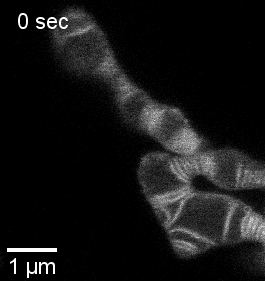
